# Social Jetlag Has Detrimental Effects on Hallmark Characteristics of Adolescent Brain Structure, Circuit Organization and Intrinsic Dynamics

**DOI:** 10.1101/2025.06.22.660935

**Authors:** Matthew Risner, Eliot S Katz, Catherine Stamoulis

## Abstract

**Study Objectives:** To investigate associations between social jetlag and developing brain circuits and structures in adolescents.

**Methods:** N = 3507 youth (median (IQR) age = 12.0 (1.1) years; 50.9% females) from the Adolescent Brain Cognitive Development (ABCD) cohort were studied. Social jetlag (adjusted for sleep debt (SJL_SC_) versus non-adjusted (SJL)), topological properties and intrinsic dynamics of resting-state networks, and morphometric characteristics were analyzed.

**Results:** Over 35% of participants had SJL_SC_ ≥2.0 h. Boys, Hispanic and Black non-Hispanic youth, and/or those at later pubertal stages had longer SJL_SC_ (β=0.06 to 0.68, CI=[0.02, 0.83], p≤0.02), which was also associated with higher BMI (β=0.13, CI=[0.08, 0.18], p<0.01). SJL_SC_ and SJL were associated with weaker thalamic projections (β=- 0.22, CI=[−0.39, −0.05], p=0.03), potentially reflecting a disrupted sleep-wake cycle. Longer SJL_SC_ was also associated with less topologically resilient and weakly connected salience network (β=-0.04, CI=[−0.08, −0.01], p=0.04), and lower thickness and/or volume of cortical and subcortical structures overlapping with this and other networks supporting emotional and reward processing and regulation, and social function (β=- 0.08 to −0.05, CI=[−0.12, −0.01], p<0.05). SJL_SC_ and SJL were associated with alterations in spontaneous brain activity and coordination that indicate disrupted neural maturation and plasticity. SJL was associated with lower information transfer between regions supporting sensorimotor integration, social function and emotion regulation (β=-0.07 to-0.05, CI=[−0.12, −0.01], p<0.04).

**Conclusions:** Misaligned sleep may have detrimental effects on adolescent brain circuit organization and dynamics, and structural characteristics of regions that play critical roles in cognitive function and regulation of fundamental biological processes.

## 1. INTRODUCTION

Adolescence is characterized by profound changes in sleep duration, state distribution, and timing, as a result of maturating regulatory processes and social environmental factors. [1] Adolescents often sleep less than recommended for optimal development, obtain lower-quality sleep, and experience substantial reductions in slow wave sleep. [1–4] Additionally, in adolescents a mismatch emerges in sleep timing between school and free days, resulting in misalignment between their biological clock and social schedule, termed social jetlag. [5]

Social jetlag is defined as a greater than 2-hour disparity in sleep timing between school days and free days. Almost half of all adolescents experience social jetlag, which has been linked to adverse physical, cognitive, and mental health outcomes in youth. Prior studies have reported associations, especially in girls, with elevated BMI and weight gain over time, [6–8] likely resulting from decreases in physical activity, higher screen time, and metabolic and endocrine dysregulation, [9–11] but also changes in food-related behaviors, including irregular meal timing, reduced intake of nutrient-dense foods, and elevated sugar intake, [12–14] and poor fitness. [15] Social jetlag has also been associated with sleep disturbances in youth, including poor sleep quality, insomnia, fatigue, and daytime sleepiness. [12, 16–20]

Social jetlag may also adversely impact cognitive function in youth, including worse academic performance, lower crystallized intelligence, poor reading performance, and memory and attention deficits, especially in girls. [19–28]

Common mental health disorders commonly emerge during adolescence, likely as a result of profound environmental, physiological and brain changes. Although their underlying causes are incompletely understood, [29, 30] inadequate (including misaligned) sleep may increase risk for these disorders. Social jetlag has been associated with higher risk of anxiety, depression, social dysfunction and antisocial behaviors, mood changes, conduct issues, and risky behaviors especially in girls. [19, 20, 31–37] Social jetlag has also been linked to higher screen time (both throughout the day and before sleep), which affects mental health and emotion regulation in youth. [9, 38–42]

Relatively few studies have examined the impact of social jetlag on the developing brain, especially during adolescence and young adulthood, as high-level cognitive function and its neural substrates continue to maturate. A recent study in young adults reported links between social jetlag and widespread alterations in functional connectivity of brain networks involved in attention, reward processing, and executive control. [43] Longer social jetlag was associated with weaker connections between the ventral striatum and inferior orbitofrontal cortex, and changes in connectivity between the dorsolateral prefrontal cortex (DLPFC) and frontal, parietal, and occipital lobes. Another recent study based on the large Adolescent Brain Cognitive Development (ABCD) study cohort, [44] reported inverse associations between social jetlag and resting-state connectivity between the right hippocampus and cingulo-opercular network. [27] Social jetlag has also been associated with lower gray matter volume of the medial prefrontal cortex, amygdala, and hippocampus. [45] Additional studies, focusing on the effects of circadian disruption and irregular sleep patterns have reported widespread brain alterations, especially in regions supporting emotional and sensory processing. [46, 47] These studies have also shown that the prefrontal cortex, which undergoes heightened maturation in adolescence, is particularly sensitive to changes in sleep patterns, with potentially long-term implications for emotional regulation and mental health. [48]

Despite their insights, prior studies on detrimental effects of social jetlag on cognitive development have provided limited knowledge on its neuromodulatory mechanisms. To date, there are no investigations that have examined its effects on fundamental brain characteristics that play central roles in cognitive function, including the organization of developing networks (i.e., beyond just the strength of their connections) and their spontaneous (intrinsic) dynamics, morphometric characteristics of their constituent structures, and communication between their regions, and their dynamics. Extensive prior work has linked these characteristics to developmental processes, cognitive function, and mental health in youth. [49–60] However, none of these studies have examined the impact of social jetlag, which increases with age in adolescence, [61] in parallel with neural maturation and profound reorganization of brain circuits, and may amplify the risk of miswiring, resulting in cognitive deficits and mental health issues.

To address this critical gap in knowledge, this study investigated associations between social jetlag and topological, morphometric, and dynamic brain characteristics in a sample of over 3,500 adolescents from the ABCD cohort, at the study’s two-year follow-up (ages ∼11-12 years). It leveraged available multimodal sleep and brain data, and used cutting-edge computational approaches to comprehensively characterize the topological organization of resting-state brain networks and information flow between their constituent regions, as well as regional and network intrinsic dynamics. The study tested the following hypotheses: 1) Social jetlag adversely impacts hallmark topological and structural characteristics of underdeveloped (and thus vulnerable) brain networks, including their efficiency, resilience, and strength of their connections; 2) It disrupts normative communication between brain regions and information transfer, including in hubs, i.e., highly connected, domain-general regions that receive extensive domain-specific information, synthesize and distribute the output to specially distributed areas, in response to cognitive demands; 3) It impairs the spontaneous coordination of brain networks, including the underdeveloped Default-Mode Network (DMN), which is active at rest and maturates significantly adolescence, [62] plays a ubiquitous role in cognitive function, [63, 64] and is adversely affected by irregular sleep patterns and poor quality sleep [65–67]; 4) Social jetlag also disrupts pubertal changes in temporal fluctuations of the DMN’s and other networks’ intrinsic coordination patterns and regional activity, i.e., it impairs the increased consistency of these patterns, which reflects normative development. [53]

## 2. METHODS

The study was approved by the Institutional Review Board.

### 2.1 Participants

Effects of social jetlag on the brain were investigated in a sample of n = 3507 typically developing adolescents (median age = 12.0 years, interquartile range (IQR) = 1.1 years; 50.9% girls). All participants had at least one high-quality 5-minute resting-state fMRI scan that was minimally affected by motion in the scanner. Neurodevelopmental and neuropsychiatric disorders can have significant adverse effects on sleep, and brain structure and circuitry. To minimize associated confounding effects, youth with a diagnosis of Attention Deficit Hyperactivity Disorder (ADHD), Autism Spectrum Disorder (ASD), or any neuropsychiatric disorder (including schizophrenia, bipolar disorder and any psychotic disorder) were excluded. In addition, those with identified anomalies in their structural MRI were also excluded.

### 2.2 Social Jetlag Measures

Two measures of social jetlag were derived from the youth-reported Munich Chronotype Questionnaire (MCTQ). [68] Similar to prior work, [27] the primary measure used in this study corrected for sleep debt (e.g., shorter sleep on school days versus oversleeping on free days), using the correction [69]:

*Social Jetlag (SJL)_SC_ = (SOW + 0.5*SD_week_) – (SOF + 0.5*SD_week_) = |SOW - SOF|* This measure corresponds to the absolute difference between sleep onset on school (SOW) and free (SOF) days. Results were also analyzed based on the classic social jetlag measure, which includes sleep debt (SJL), [5] i.e., the absolute difference between sleep midpoints (the time halfway between sleep onset and wake time) on school and free days, which includes weekend oversleep due to sleep debt. Both measures are expressed in hours.

### 2.3 Neuroimaging data

Structural and functional (resting-state) MRI data were collected in 3.0T scanners (GE, Siemens, Philips; repetition time (TR) = 0.8 s; isotropic voxel size = 2.4 mm) at 21 ABCD study sites across the United States. The fMRI protocol included up to four, 5-min runs, separated by short videos to ensure that participants did not fall asleep in the scanner. Prior to sharing with the community, data undergo at least minimal preprocessing at the ABCD study’s Data Analysis, Informatics & Resource Center (DAIRC) to correct for head movement, B0 distortions, and gradient nonlinearities. [70] Minimally processed fMRI and fully processed structural MRI were analyzed in this study. Details on processing fMRI data using the custom Next Generation Neural Data Analysis (NGNDA) platform are provided in Supplemental Materials and Brooks et al., 2021. [65]

#### 2.3.1. Time-independent and dynamic topological properties

Time-independent (non-directional, time-compressed), effective (directional, time-compressed) and time-dependent (dynamic) connectivity matrices were estimated using approaches developed and/or applied in our prior work [53, 65]. Details on their estimation are provided in Supplemental Materials. Topological properties from time-compressed and dynamic connectivity matrices were estimated at three spatial scales: a) the entire brain (connectome), b) large-scale resting-state networks, [71] and additional ones, including the reward, [72] prefrontal cortical (and its projections, as separate circuits), social, [73] and central executive networks, [74] and c) individual brain regions (network nodes). These properties included measures of connection strength, community structure, topological resilience, efficiency, fragility, and regional importance in the network, and are listed in Supplemental Materials.

#### 2.3.2 Temporal variability of spontaneous brain activity and coordination

Time-varying connectivity and corresponding adjacency matrices were estimated from fMRI signals using a sliding window approach (with a 16.0s (20 frames) window length, and 1-frame sliding). The estimation is described in detail in Supplemental Materials and Lim et al., 2025. [53] The coefficient of dispersion was selected as the estimator of topological property variability over time.

Temporal fluctuations of regional activity were also estimated. Following the approach in Sydnor et al., 2023, [75] the frequency power spectrum of each parcel signal was estimated, and the median square root of low-frequency spectral power (0.01 - 0.10 Hz) was used to quantify signal amplitude fluctuations. Then, parcel-level estimates were downsampled to 100 regional fluctuation amplitudes by taking the median over all parcels within a region. [53]

#### 2.3.3 Information transfer between brain regions

To estimate information flow to and out of a region, effective connectivity (directional interactions) between pairs of regions was estimated. [76, 77] Phase transfer entropy (PTE) [78] was selected as the estimator of this measure, and is described in detail in Supplemental Materials. For each brain region, median information inflow, outflow and net flow were estimated.

#### 2.3.4 Morphological brain properties

Cortical thickness, white matter intensity (potentially reflecting developmental white matter differences), and cortical and subcortical grey matter volume were estimated from fully processed T1-weighted MRI scans, and provided by the ABCD. These morphometric measures were estimated in 68 cortical and 30 subcortical and cerebellar regions. Two atlases were used for parcellation, the Desikan-Killiany atlas [79] for the cortex, and one based on probabilistic classification, [80] for subcortical regions and the cerebellum.

### 2.4 Statistical analysis

Linear mixed-effect models that included random effects (intercept and slope) for ABCD site, to account for potential site effects, were developed for all analyses. In addition, these sites are geographically diverse and vary substantially as a function of population density. Thus, all analyses also accounted for sampling bias using propensity scores provided by the ABCD study. Age, sex, pubertal stage, race-ethnicity, family income, and BMI (z-scores stratified by sex) [81] were included in all models. Due to insufficient statistical power for comparisons of racial groups, race-ethnicity was combined into a binary variable: white non-Hispanic (0) versus racioethnic minority (1). Pubertal stage ranged from pre-puberty (1) to late/post-puberty (4). Models that included brain parameters were also adjusted for time of day of the scan [82] and percent of frames censored for motion. In these models, SJL_SC_ or SJL were the independent variable of interest, and brain characteristics the dependent variables. Additional models assessed relationships between social jetlag and demographic and other individual characteristics, as well as total weekly screen time (in hours). In these models, SJL_SC_ or SJL were the dependent variable. The statistical significance level was set at α = 0.05. Given multiple comparisons (for example, multiple topological or morphometric properties, all p-values were corrected for the False Discovery Rate (FDR) [83]. At the connectome, network levels and structural region levels, corrections were done over all topological (or morphometric) properties of the entire brain, a network or a region. At the node/region level, p-values were corrected over all nodes in a particular network.

Model validation used predictive power as the relevant metric. The sample was randomly split into training and testing sets (75:25), and model parameters were estimated from the training set. The process was repeated 100 times, and at each iteration the Coefficient of Variation of the Root Mean Square Error (CV[RMSE]) was estimated (using the testing set) as a measure of the model’s predictive power. Models with CV[RMSE] ≤0.2 were considered to have good predictive power.

## 3. RESULTS

Almost 40% of participants were in pre or early puberty (n = 1383 (39.4%)), and ∼30% in mid-puberty (n = 1133 (32.3%)). The distribution of race and ethnicity, and family income of the sample reflected that of the ABCD study. Over 50% of participants were white and non-Hispanic (n = 1820 (51.9%)), ∼15 were black and non-Hispanic (n = 507 (14.5%), and ∼20% were Hispanic (n = 771 (22.0%)). About 47% of families (n = 1646 (46.6%)) had a yearly household income of <$100,000. Median BMI was 19.3 kg/m^2^ (IQR = 5.6). Participant characteristics are summarized in Table 1.

**Table 1.** Participant characteristics (n = 3507). The racioethnic category “Other Non-Hispanic” combines small racial categories: Filipino, Vietnamese, Alaska Native, American Indian, Asian Indian, Chinese, Guamanian, Hawaiian, Japanese, Korean, Native Samoan, other Pacific Islander, other Asian, multiracial, other race.

| Characteristic | Statistic/Category | Value |
| --- | --- | --- |
| Age (years) | Median (IQR) | 12.00 (1.08) |
| Sex | Male | 1720 (49.04%) |
|  | Female | 1785 (50.90) |
|  | Missing | 2 (0.06%) |
| Race/Ethnicity | White Non-Hispanic | 1820 (51.90%) |
|  | Black Non-Hispanic | 507 (14.46%) |
|  | Asian Non-Hispanic | 68 (1.94%) |
|  | Other Non-Hispanic | 330 (9.41%) |
|  | Hispanic | 771 (21.98%) |
|  | Missing | 11 (0.31%) |
| Family Income (\$) | <25,000 | 344 (9.81%) |
|  | 25,000 - 49,999 | 416 (11.86%) |
|  | 50,000 - 74,999 | 426 (12.15%) |
|  | 75,000 - 99,999 | 450 (12.83%) |
|  | 100,000 - 199,999 | 1094 (31.19%) |
|  | ≥200,000 | 508 (14.49%) |
|  | Missing | 269 (7.67%) |
| BMI | Median (IQR) | 19.33 (5.58) |
|  | Missing | 27 (0.77%) |
| Pubertal Stage | Pre-Puberty | 634 (18.08%) |
|  | Early Puberty | 749 (21.36%) |
|  | Mid-Puberty | 1133 (32.30%) |
|  | Late/Post-Puberty | 801 (22.84%) |
|  | Missing | 190 (5.42%) |

Median (IQR) SJL_SC_ was 1 (1) h (and 1.5 (1.5) h based on SJL ). About 74% of the sample had SJL_SC_ ≥1 hour, and ∼36% had ≥2 hours. Girls, white non-Hispanic youth, and those from families with higher family income had shorter SJL_SC_ (β = −0.11, CI = [- 0.23, −0.0003], p = 0.05; β = −0.45, CI = [−0.55, −0.35], p < 0.01; β = −0.11, CI = [−0.13, −0.08], p < 0.01, respectively). In contrast, Hispanic and Black non-Hispanic youth had longer SJL_SC_ (β = 0.14, CI = [0.02, 0.25], p = 0.02; β = 0.68, CI = [0.53, 0.83], p < 0.01), and so did those at later pubertal stages (β = 0.06, CI = [0.01, 0.12], p = 0.03, respectively). BMI was positively associated with SJL_SC_ (β = 0.13, CI = [0.08, 0.18], p < 0.01), and similarly for total weekly screen time (β = 0.008, CI = [0.006, 0.011], p < 0.01). Model statistics are provided in Table 2.

**Table 2.** Statistics of models testing associations between social jetlag excluding sleep debt (SJL_SC_) and demographic and other characteristicsAll p-values have been adjusted for the False Discovery Rate. CI: Confidence interval.

| Covariate | Beta | 95th % CI | P-value |
| --- | --- | --- | --- |
| Pubertal stage | 0.063 | [0.006, 0.121] | 0.032 |
| Sex | -0.113 | [-0.227, -0.001]] | 0.049 |
| White non-Hispanic | -0.446 | [-0.546, -0.345] | <0.001 |
| Black non-Hispanic | 0.680 | [0.528, 0.833] | <0.001 |
| Hispanic | 0.135 | [0.023, 0.248] | 0.018 |
| Family income | -0.107 | [-0.133, -0.082] | <0.001 |
| BMI | 0.127 | [0.078, 0.176] | <0.001 |
| Screen time | 0.008 | [0.006, 0.011] | <0.001 |

### 3.1 Associations between social jetlag and time-independent topological characteristics

#### 3.1.1 Network topological properties

Longer SJL_SC_ was associated with lower median connectivity between the right thalamus and the rest of the brain (β = −0.22, CI = [−0.39, −0.05], p = 0.03). It was also associated with lower median within-network connectivity (β = −0.04, CI = [−0.08, −0.01], p = 0.04), and higher fragility (β = 0.05, CI = [0.01, 0.09], p = 0.04) of the right salience network. In addition, longer SJL was associated with lower median connectivity (within- and across-network) of the right thalamus (β = −0.29, CI = [−0.48, −0.10], p < 0.01), lower efficiency and lower median connectivity between the central visual network and the rest of the brain (β = −0.05, CI = [−0.10, −0.01], p < 0.04), and lower efficiency of the right (β = −0.05, CI = [−0.10, −0.01], p = 0.04) and higher fragility of the bilateral peripheral visual network (β = 0.06 to 0.08, CI = [0.01, 0.15], p < 0.05). Model statistics are provided in Table 3. All models had good predictive power (CV[RMSE] < 0.20)).

**Table 3.**
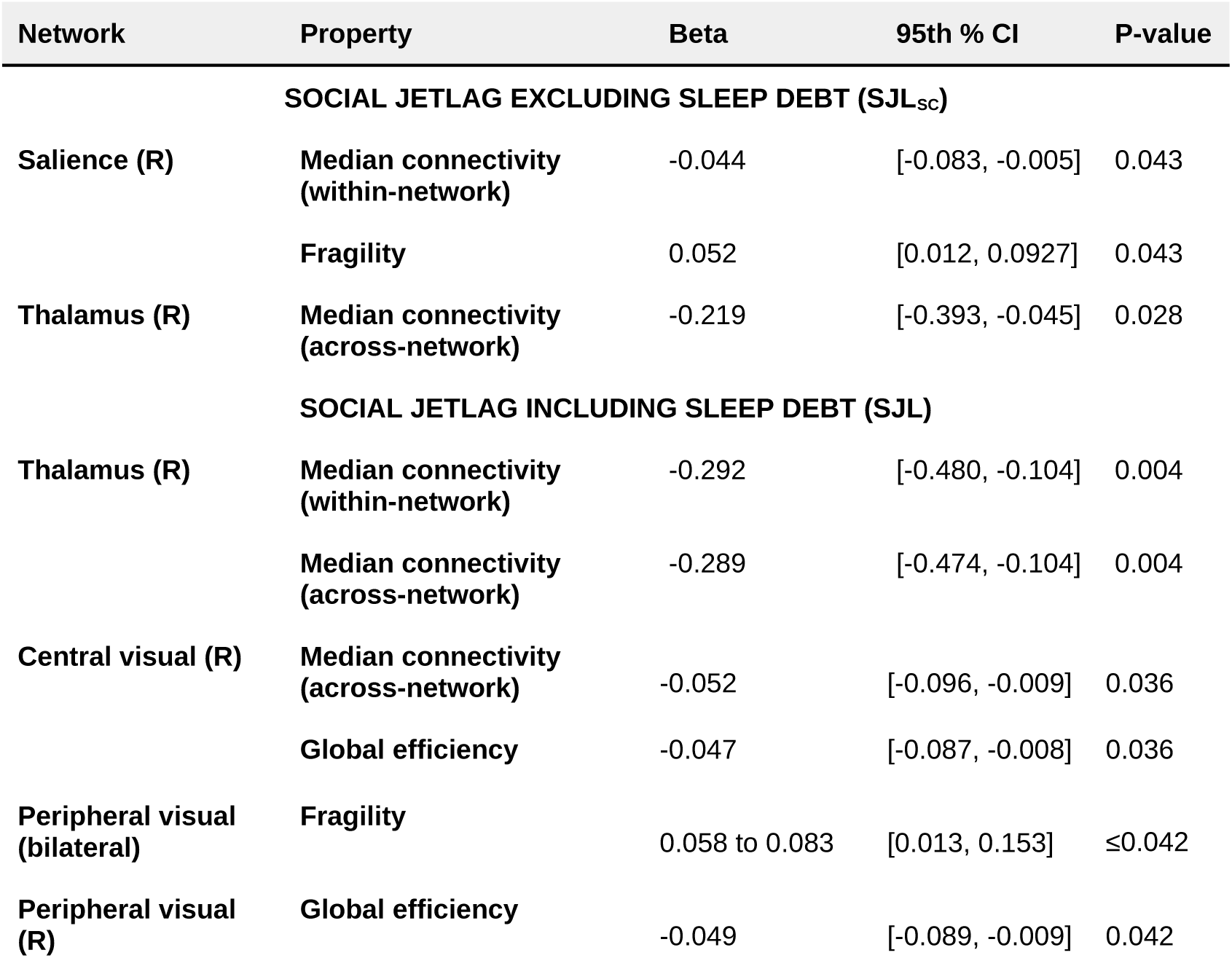
Statistics of models testing associations between SJL_SC_ (excluding sleep debt) and topological properties of individual networks, and similarly for SJL (including sleep debt) Results are based on the best-quality run. All p-values have been adjusted for the False Discovery Rate. CI: Confidence interval.

#### 3.1.2 Local (regional) topological properties

At the parcel (node) level, and based on the best run, SJL was associated with lower local clustering within the bilateral central and peripheral visual networks (β = −0.09 to - 0.05, CI = [−0.13, −0.01], p < 0.05), higher centrality of the left somatomotor network (β = 0.05 to 0.08, CI = [0.01, 0.12], p < 0.05) and lower centrality of the bilateral dorsal attention and central and peripheral visual networks (β = −0.09 to −0.05, CI = [−0.13, - 0.01], p < 0.05). These associations are shown in Figure 1. There were no corresponding statistical associations with SJL_SC_.

**Figure 1.**
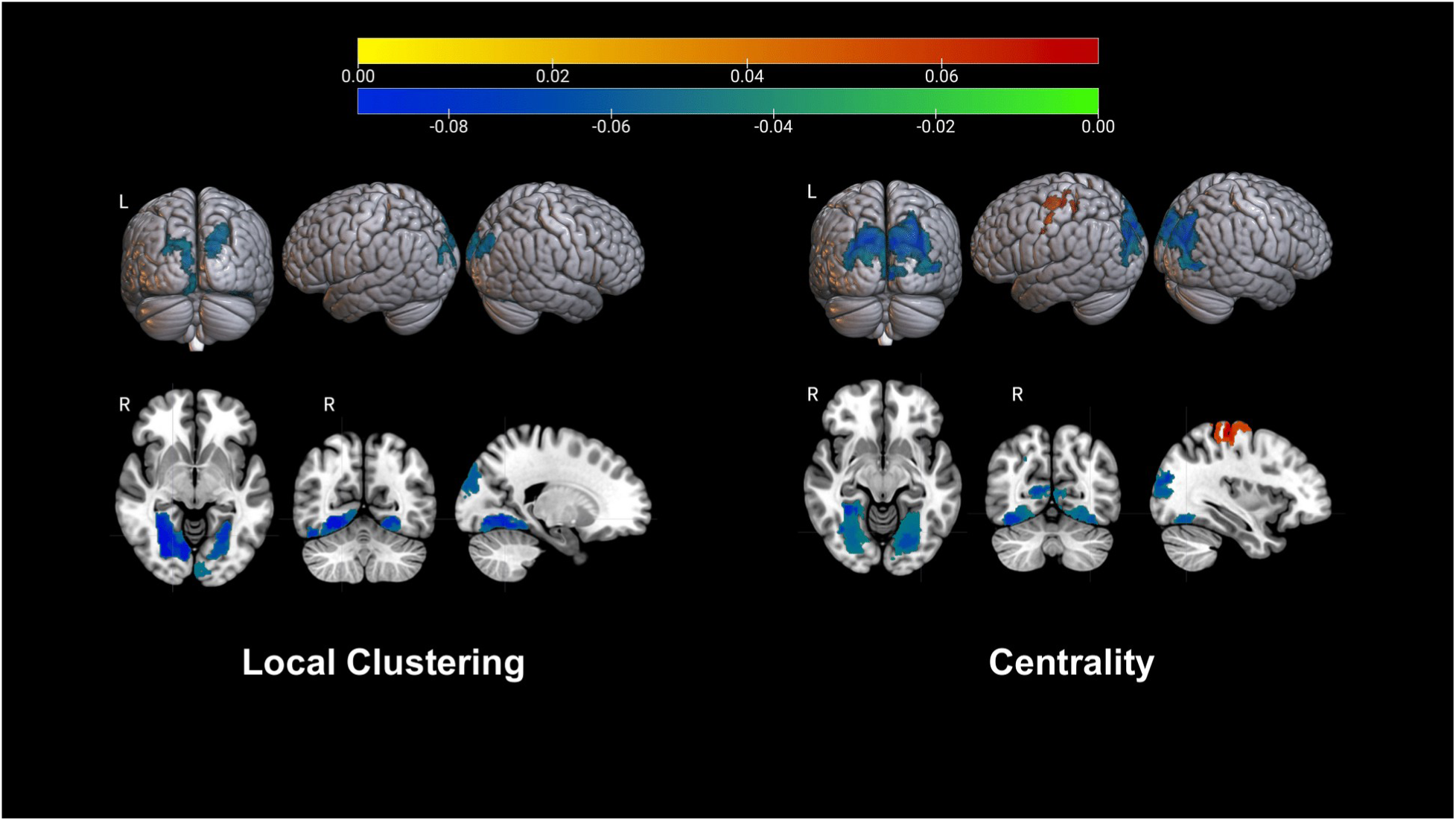
Associations between SJL (including sleep debt) and local clustering (left panels) and node centrality (right panels). The color map represents the values of model regression coefficients. Yellow to red corresponds to positive values and blue to green to negative values.

### 3.2 Associations between social jetlag and morphometric regional properties

SJL_SC_ was associated with lower thickness of multiple structures of the temporal lobe (including the banks of the left superior temporal sulcus, bilateral superior temporal gyrus and right medial temporal gyrus) and the bilateral lingual gyrus (β = −0.07 to −0.05, CI = [−0.12, −0.01], p < 0.04). It was also associated with lower volume of the banks of the left superior temporal sulcus, bilateral middle and left inferior temporal gyri, left entorhinal cortex, bilateral inferior parietal gyrus, left insula, left hippocampus, and bilateral caudate nucleus, putamen, amygdala, and nucleus accumbens (β = −0.08 to - 0.05, CI = [−0.12, −0.01], p < 0.05). These associations were overall consistent for SJL as well. Additional negative associations were estimated between SJL and volume of multiple regions, including lateral and medial orbitofrontal cortces and several brain hubs, such as the bilaeral precuneus and the right cingulate cortex. All models had good predictive power (CV[RMSE] < 0.20). Model statistics are provided in Table 4.

**Table 4.**
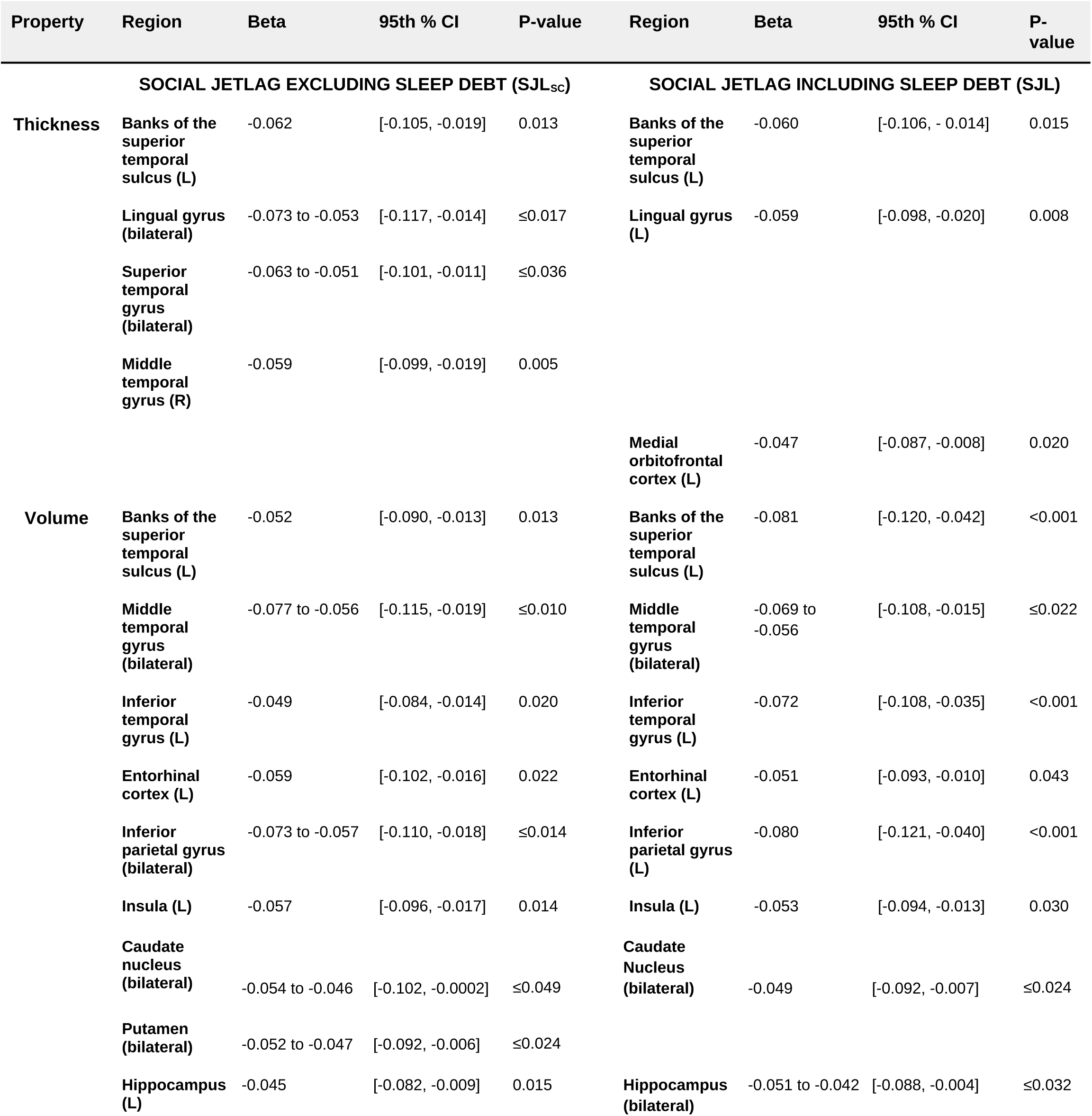
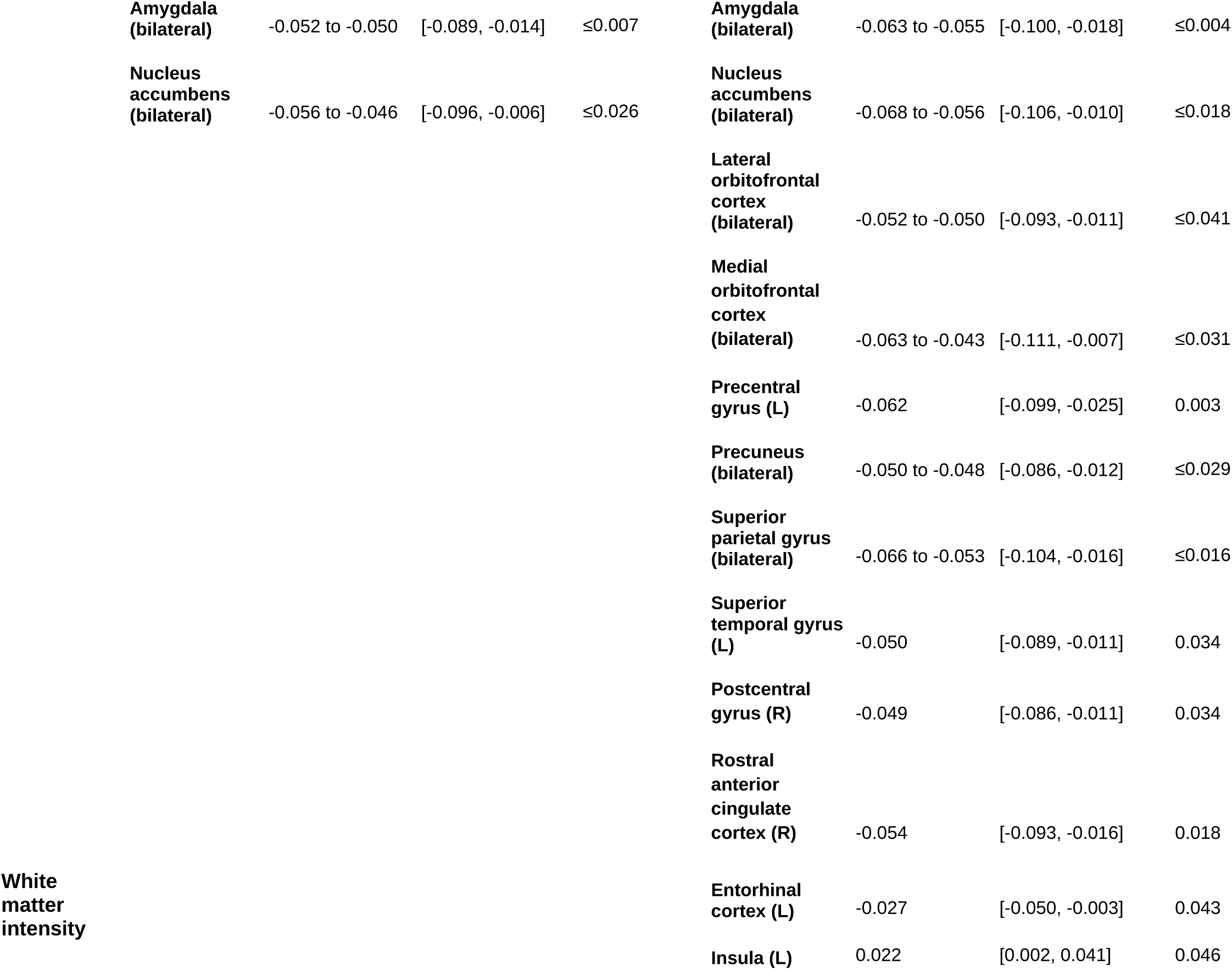
Statistics of models testing associations between SJL_SC_ (excluding sleep debt)and morphometric brain properties, and similarly for SJL (including sleep debt). All p-values have been adjusted for the False Discovery Rate. CI: Confidence interval.

### 3.3 Associations between social jetlag and signal and topological dynamics

SJL_sc_ was associated with higher local clustering fluctuations in bilateral somatomotor areas (β = 0.05, CI = [0.01, 0.10], p < 0.04). SJL was also associated with higher clustering fluctuations in a left somatomotor region (β = 0.06, CI = [0.02, 0.10], p = 0.02) and lower clustering fluctuations in the salience, dorsal attention, and basal ganglia regions (β = −0.07 to −0.05, CI = [−0.12, −0.01], p < 0.03). The spatial distributions of these associations are shown in Figure 2. In addition, SJL_SC_ was associated with higher fluctuation amplitude in left somatomotor areas (β = 0.04 to 0.05, CI = [0.004, 0.09], p < 0.04) and lower fluctuation amplitude in bilateral dorsal attention, left frontoparietal control, and left default mode areas (β = −0.06 to −0.04, CI = [−0.11, −0.01], p < 0.05). SJL was also associated with higher fluctuation amplitude of bilateral somatomotor areas (β = 0.05 to 0.08, CI = [0.01, 0.12], p < 0.05) and lower fluctuation amplitude of bilateral dorsal attention, bilateral frontoparietal control, bilateral default mode, and right salience areas (β = −0.08 to −0.05, CI = [−0.13, −0.01], p < 0.05). All related statistical models had good predictive power (CV[RMSE] < 0.13). The spatial distribution of these associations is shown in Figure 3.

**Figure 2.**
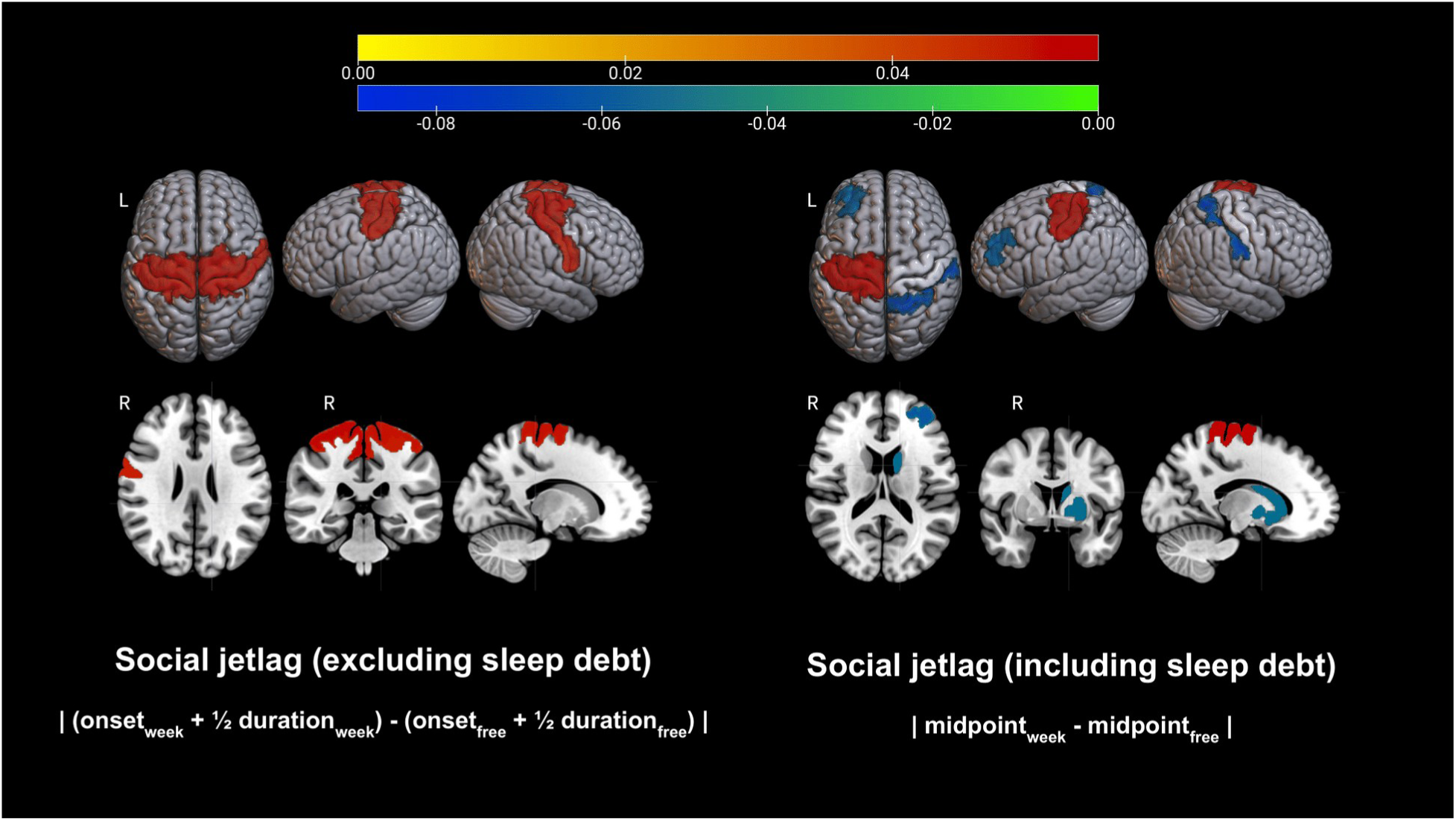
Associations between social jetlag and local clustering fluctuation. The color map represents the values of model regression coefficients. Yellow to red correspond to positive values and blue to green to negative values.

**Figure 3.**
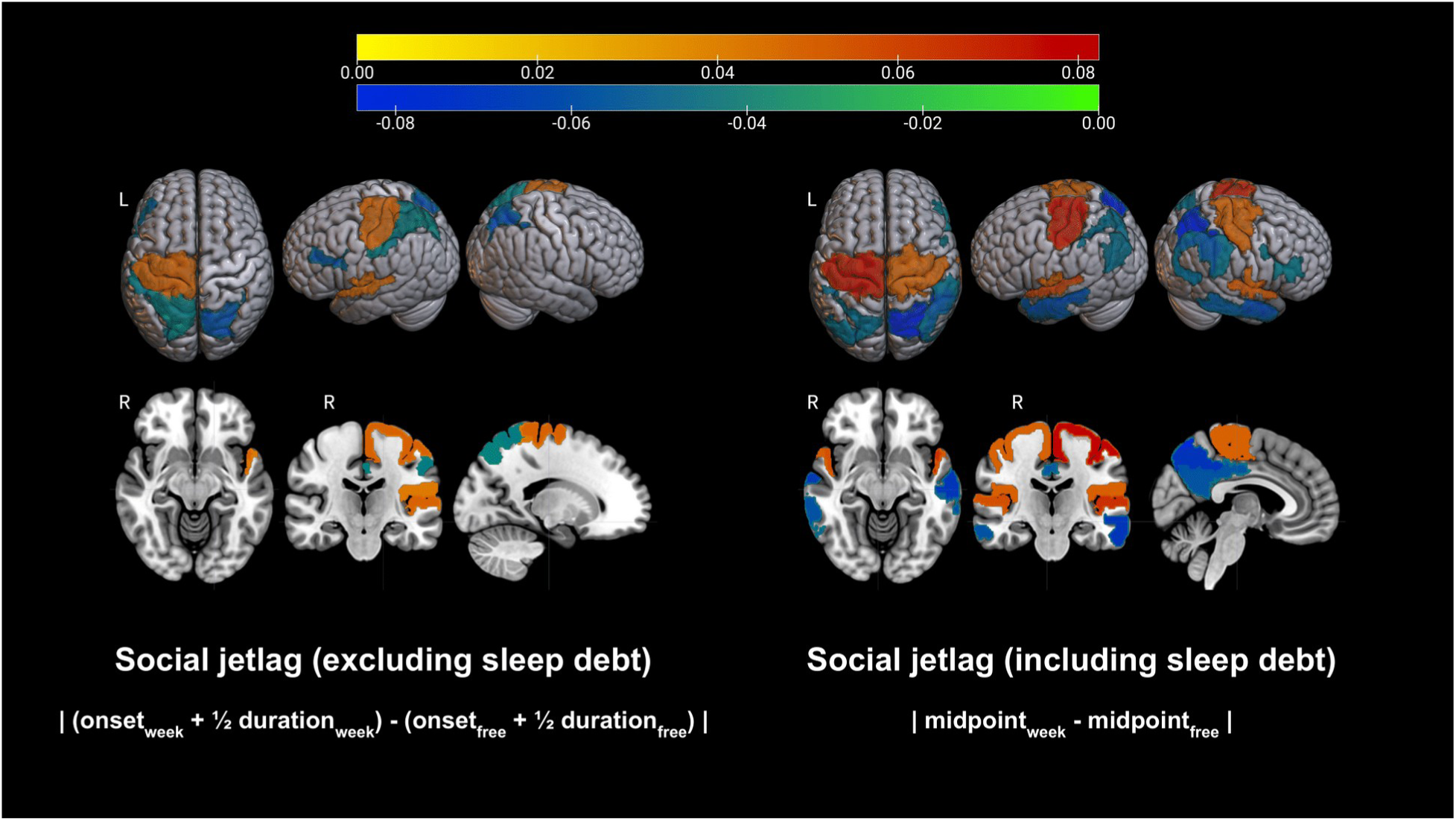
Associations between SJL_SC_ (excluding sleep debt) – left panels, and SJL (including sleep debt) – right panels, and fluctuation amplitude. The color map represents the value of model regression coefficients. Yellow to red corresponds to positive values and blue to green to negative values.

### 3.4 Associations between social jetlag and regional information flow

SJL was associated with lower median outflow from the left amygdala, bilateral temporoparietal, right peripheral visual, and right somatomotor regions (β = −0.07 to - 0.04, CI = [−0.11, −0.003], p < 0.05). It was also associated with lower median net flow in the right temporoparietal, peripheral visual, and somatomotor areas (β = −0.07 to −0.05, CI = [−0.12, −0.01], p < 0.04). All related models had good predictive power (CV[RMSE] < 0.09). The spatial distributions of these associations are shown in Figure 4. No corresponding associations were estimated for SJL_SC_.

**Figure 4.**
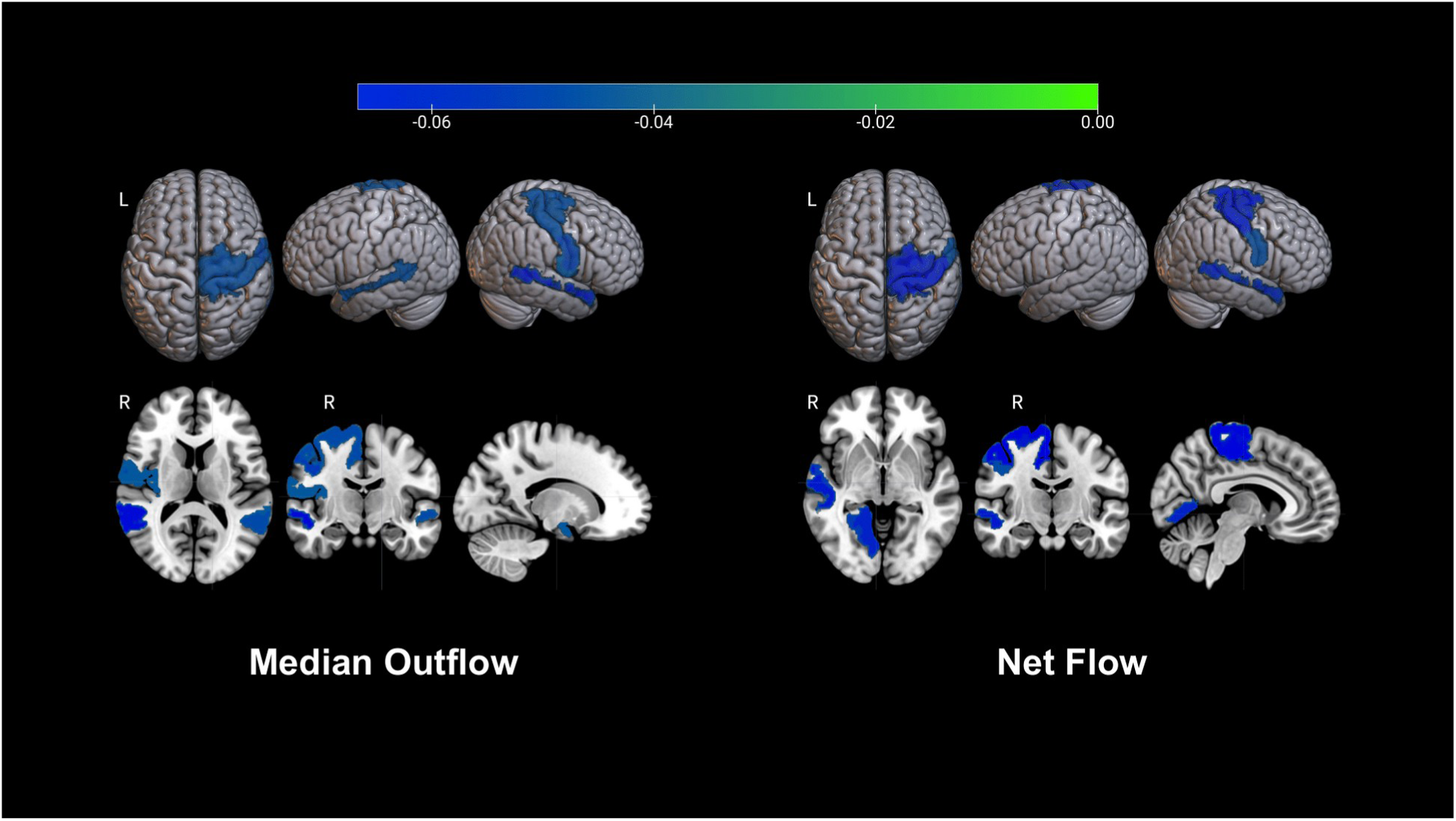
Associations between SJL (including sleep debt) and information flow. The color map represents the value of model regression coefficients (which were all negative).

### 3.5 Mappings between structural and functional correlates of social jetlag

Connectivity between the right central visual network and the rest of the brain, and morphometric properties of the right lingual gyrus (which overlaps with this network) were positively associated (β = 0.15, CI = [0.09, 0.21], p < 0.01), and both were negatively associated with SJL_SC_. Topological fragility of the bilateral visual peripheral network was negatively associated with morphometric properties of the bilateral lingual gyrus (β = −0.12 to −0.11, CI = [−0.17, −0.06], p < 0.01), and SJL_SC_ was associated with higher fragility of the network and lower thickness of the gyrus. Furthermore, fragility of the bilateral visual peripheral network was negatively associated with morphometric properties of the overlapping bilateral superior parietal gyrus (β = −0.11 to −0.09, CI = [- 0.15, −0.04], p < 0.01), and SJL was associated with higher fragility of the network and lower volume of the gyrus. Finally, higher fluctuation amplitude in the right somatomotor network was negatively associated with out and net flow in its constituent regions, (β = - 0.22 to −0.12, CI = [−0.26, −0.07], p < 0.01), with both properties being negatively impacted by SJL_SC_ (higher fluctuation amplitude and lower information flow). Statistics on additional associations are provided in Table S1.

## 4. DISCUSSION

Insufficient, poor quality and/or misaligned sleep during development has profound and often long-lasting detrimental effects on brain health. During sensitive periods, such as adolescence, these effects may be amplified, leading to cognitive deficits (including impaired learning) and mental health issues. Although the effects of insufficient and disrupted/disordered sleep on cognitive and mental health have been extensively studied, those of misaligned sleep (and social jetlag) remain incompletely understood. Prior research, including large epidemiological studies, has shown that social jetlag affects over half of adolescents and adversely impacts their academic performance, high-level cognitive function (including executive control), emotional regulation, and mental health. However, its mechanism of action on underlying brain circuits and hallmark characteristics of their organization, morphoogy and activity are elusive. Recent work on social jetlag based on the ABCD cohort, has focused only on functional connectivity, and thus connection strength [27]. However, cognitive function critically depends on the organization of brain circuits, and properties of their constituent structures.

In this study we have addressed this gap in knowledge and, in a sample of over 3,500 adolescents, have investigated associations between social jetlag and comprehensive brain properties across multiple domains, including topology, morphology, and dynamics. It examined and compared both SJL_SC_ and SJL, i.e., social jetlag excluding sleep debt (to minimize effects of insufficient sleep on school days and oversleeping on free days), and social jetlag that included sleep debt. To the best of our knowledge this is the first study to examine impacts of social jetlag on the evolving organization and dynamics of adolescent resting-state networks - the backbone of the functional connectome, properties of their constituent structures, and information flow through the brain, which is critical to information processing and cognitive function.

Over a third of participants had social jetlag of at least 2 h, an alarming statistic given its links to increased risk for mental health problems, [84] cardiovascular, metabolic, and hormonal issues, and obesity [6, 85–87]. Longer social jetlag was also associated with longer weekly screen time, as well as BMI, in agreement with prior studies that have reported associations between screen time and social jetlag, [40] a sedentary lifestyle, lower physical activity and higher BMI [9, 88]. In addition, Black and Hispanic youth had longer social jetlag, again a finding that is aligned with a number of prior studies that have reported significant sleep disparities, including misaligned and poor quality sleep in racial and ethnic minorities [89–92].

Equally alarming are the identified associations between social jetlag and fundamental aspects of brain structure, organization of its circuitry, and intrinsic dynamics, all of which play critical roles in cognitive function. Longer social jetlag (both excluding and including sleep debt) was associated with weaker and/or less resilient (more topologically fragile) connections between the thalamus and the rest of the brain. In addition, SJL was associated with weaker, less efficient and/or more fragil connections in visual networks, while SJL_SC_ was associated with weaker connections within the salience network.The thalamus plays a critical role in the regulation of the sleep-wake cycle and circadian rhythm, which may be disrupted by social jetlag. [93, 94] Our results suggest a mechanistic relationship between social jetlag and the regulation of the sleep-wake cycle, through the former’s impact on the thalamus. In addition, prior studies have associated social jetlag with alterations in brain networks that include the thalamus, and support motor function and posture control, but are also involved in reward processing, emotion regulation and eating behaviors, as well as mental health [32, 95, 96]. Our findings suggest another potential mechanistic relationship between social jetlag and these processes, through its adverse effects on networks that regulate them and/or support mental health.

Social jetlag was also associated with lower fluctuations of spontaneous brain activity and coordination patterns in some regions, and higher fluctuations in others. Prior work, including our own, has shown that intrinsic topological and BOLD signal fluctuations represent markers of brain development, and broadly decrease with age and neuroanatomical maturation, i.e., spontaneous coordination patterns become increasingly consistent. [53, 75] Prior studies have also shown that higher spontaneous BOLD fluctuations in sensorimotor areas may be associated with impaired plasticity [97]. Both social jetlag measures were associated with higher signal and topological fluctuations in these areas, suggesting potential detrimental effects of social jetlag on fundamental processes that facilitate adaptation and learning. During development, changes in intrinsic brain dynamics are spatially heterogeneous, and may be disrupted by social jetlag, likely through its adverse effects on underlying structures, resulting in less consistent coordination patterns and higher signal variability in some areas, and abnormally lower fluctuations in others. This potential mechanism of action is partially supported by identified negative associations between social jetlag and cortical and subcortical volume of spatially distributed brain areas, overlapping with regions and networks in which topological and dynamic properties were adversely impacted by social jetlag. These included structures of the salience network, including the insula, amygdala and the nucleus accumbens, which play central roles in reward and emotion processing, and regions of the DMN, attention and frontoparietal networks. Together, these support cognitive processes that develop significantly in adolescence, including executive control and decision-making but also reward and emotion regulation [98–101]. Thus, social jetlag may have detrimental effects on these processes through its neuromodulatory effects on brain structures, circuits and their intrinsic dynamics.

We have also identified distributed negative associations between social jetlag that included sleep debt and information flow through the brain, including hubs, which receive, synthesize and output information in support of cognitive demands. Communication between brain regions is critical to cognitive processing. Some domain-specific regions may receive more information (for example external (sensory) inputs) than they output, and others may output more information than they receive, with cognitive and/or topological hubs often receiving and outputting equal amounts of information [102–103]. Social jetlag was associated with lower flow of information from the amygdala to the rest of the brain, and similarly from temporoparietal, peripheral visual, and right somatomotor regions. This implies that it may impair distributed interactions between brain regions that support sensory processing, motor function, sensorimotor interactions, but also high-level social function and emotional processing. These associations were specific to social jetlag that included sleep debt, and may thus be partly resulting from it. Nevertheless, they were specific to regions where other negative associations were identified for both measures of social jetlag. Together with its topological, structural, and dynamic correlates, these findings suggest that social jetlag may have extensive detrimental effects on the adolescent brain, affecting the strength, resilience and the consistency of spontaneous coordination patterns of its circuits, information transmission through them, and their underlying structures. Given the vulnerability of the brain during this sensitive period, these findings have important implications for both mental health and cognitive function across domains, and academic performance, as highlighted in prior work on another sample from the ABCD study cohort [24].

Despite its many strengths, including the large sample, and comparisons of measures of social jetlag with and without sleep debt with comprehensive measures of brain structure, topology, and dynamics, the study also had some limitations. First, social jetlag was estimated from the MCTQ, which is a subjective instrument, and thus less accurate than actigraphy-based estimates. Although the ABCD study is collecting actigraphy data, they are only available for a substantially smaller sample at the two-year follow-up, which may limit the statistical power of some analyses. Nevertheless, a future investigation could compare subjective and objective measures of social jetlag and its brain correlates in this smaller sample, similarly to Yang et al., 2023 [27]. In addition, this study was retrospective, and was thus inherently limited by the decisions made by the ABCD investigators. However, the ABCD is the only adolescent study to collect both sleep (and related measures) and multimodal neuroimaging data from such a large cohort. Thus, leveraging these data provides unique opportunities to study impacts of sleep misalignment on brain development and generate findings that may be generalizable to the larger adolescent population.

This study makes a significant contribution to the field’s incomplete understanding of the neural correlates of social jetlag in the adolescent brain. It provides novel mechanistic insights into how misaligned sleep, which is common in adolescence (almost 40% of youth in this study experienced social jetlag of at least 2 h), may impact fundamental biological processes, such as the sleep-wake cycle, through its impact on the thalamus and its connections with the rest of the brain, and evolving cognitive functions, such as reward processing and emotional regulation, through its impact on the organization and strength of brain networks that support them. Social jetlag may also modulate the brain’s task-independent (intrinsic) dynamics, and spontaneous coordination of its regions, a process that plays a ubiquitous role in cognitive function and is impaired in mental health disorders, and may disrupt developmental changes in these dynamics. Finally, social jetlag may also impair information transfer between brain regions that together support sensory processing, sensorimotor integration, emotional regulation and social function.

## Acknowledgments

This study was funded by the National Science Foundation, grants #2207733, 2116707.

## Data Availability Statement

The data underlying this article are publicly available through the National Institute of Mental Health, National Data Archive (NDA): https://nda.nih.gov/ Computer codes associated with the analyses are available at: https://github.com/cstamoulis1/Social-Jetlag-Brain

