## Supplemental materials for "Social Jetlag Has Detrimental Effects on Hallmark Characteristics of Adolescent Brain Structure, Circuit Organization and Intrinsic Dynamics"

**1. Resting-state fMRI processing**

The custom Next Generation Neural Data Analysis (NGNDA) platform was used to further process and harmonize the data across scanners (*Next-Generation Neural Data Analysis (NGNDA)*, 2020). Algorithms in the pipeline were used to further correct for head motion and exclude frames with displacement ≥0.3 mm, and suppress motion and cardiorespiratory artifacts, as well as harmonize the data across scanners. Each participant’s fMRI scans were coregistered to their structural MRI and normalized to the standard MNI152 space. Following bandpass filtering, voxel-level time series (in the frequency 0.01–0.25 Hz) were parcellated into 1088 distinct parcel signals using a combination of cortical, subcortical, and cerebellar brain atlases (Diedrichsen et al., 2009; Schaefer et al., 2018; Tian et al., 2020). Parcel level signals were further denoised to suppress residual artifacts (Brooks, et al., 2021). Primary analyses focused on the highest-quality fMRI run, selected based on the lowest median connectivity (given that the brain at rest is weakly coordinated) and typically coinciding with the run with the lowest percent of frames censored for motion (median (IQR) = 1.1 (4.0)%). For participants with more than one high-quality run, a second run was also analyzed for replication purposes (n = 2687; ~77% of the cohort; median (IQR) percent of frames censored for motion = 1.3 (3.7)%).

**2. Estimation of time-independent, effective and dynamic topological properties**

**A. Time-independent connectivity matrices and topological properties:** These matrices were estimation using peak cross-correlation between pairs of fMRI signals as a measure of connection strength (connectivity). Parcel-level weighted adjacency matrices were then estimated by thresholding corresponding connectivity matrices, using a population-level threshold derived from cohort-wide statistics. To eliminate spurious connections, a conservative threshold, equal to the moderate outlying peak cross-correlation value (median + 1.5*IQR), estimated by bootstrapping the statistic (1000 draws with replacement).

Topological properties were estimated at the whole-brain (connectome), network, and regional (node) scales. They included modularity (community structure), global efficiency and clustering, topological robustness, stability and fragility, segregation (ratio of within- to across-community connections), and median connectivity (within- and across-network). At the node level, properties included degree, local clustering, and eigenvector centrality (a measure of a region’s importance in the network. All measures were computed using algorithms implemented in the NGNDA platform.

**B. Time-dependent connectivity matrices and topological properties:** Time-varying connectivity and corresponding adjacency matrices were estimated from fMRI signals using a sliding window approach (with a 16.0s (20 frames) window length, and 1-frame sliding). In each window, a covariance matrix was calculated and transformed to a correlation matrix. To facilitate computationally tractable analyses, 1088 X 1088 parcel-level correlation matrices were downsampled to 100 X 100 region-level matrices, based on anatomically defined cortical, subcortical, and cerebellar regions. Subject-specific thresholds were estimated at this spatial resolution, and were used to derive weighted adjacency matrices (Lim et al., 2025). Across the three spatial scales, topological properties were estimated at each time point, and their variability over time was quantified using the coefficient of dispersion, a non-parametric measure given the non-normal distribution of time-dependent properties, which is equal to the difference between the 3rd and 1st quartiles, divided by their sum.

**C. Effective connectivity and information flow**

To estimate information flow into and out of a region, effective connectivity was estimated (Friston, 1994, 2011). Effective connectivity measures the influence of one region’s activity on another, and thus information transfer between them. In this study, phase transfer entropy (PTE; Lobier et al., 2013) was used to estimate this measure, at the scale of 100 regions. PTE is calculated from the instantaneous phase of pairs of signals (i,j), based on which transfer entropy is estimated, to assess the impact of the phase of signal i at time t on the phase of signal j at time t+1. PTE was estimated using an existing implementation (Fraschini & Hillebrand, 2017; Lobier et al., 2013). A time delay of three time points (~2.5 s) was assumed in the estimation. For each region, three measures were estimated: a) median flow of information out of a region (median taken over all values across each row of the PTE matrix); b) median flow of information into a region (median taken over all values across each column of the PTE matrix); c) net flow as the difference between outflow and inflow. In addition, directed PTE (dPTE), was calculated by normalizing the PTE values to the range 0-1, to reflect the preferential direction of information flow (Hillebrand et al., 2016). Values >0.5 indicated higher outflow, and those <0.5 higher inflow.

**Table S1.** Statistics of models testing associations between topological properties or fluctuation amplitude and information flow. All measures had significant associations with classic social jetlag. All p-values have been adjusted for the False Discovery Rate. CI: Confidence interval.

| **Network** | **Property** | **ROI** | **Information Flow Measure** | **Beta** | **95th % CI** | **P-value** |
| --- | --- | --- | --- | --- | --- | --- |
| **Time-compressed topological properties** | | | | | | |
| **Peripheral visual (R)** | **Global efficiency** | **Peripheral visual (R)** | **dPTE** | -0.310 | [-0.345, -0.276] | <0.001 |
|  |  |  | **PTE (out)** | -0.439 | [-0.471, -0.407] | <0.001 |
|  | **Fragility** | **Peripheral visual (R)** | **dPTE** | 0.140 | [0.105, 0.176] | <0.001 |
| **Fluctuation amplitude** | | | | | | |
| **Somatomotor (R)** | **Fluctuation amplitude** | **Somatomotor (R)** | **dPTE** | -0.117 | [-0.163, -0.070] | <0.001 |
|  |  |  | **PTE (out)** | -0.217 | [-0.261, -0.174] | <0.001 |
